# ImmuNet: A Segmentation-Free Machine Learning Pipeline for Immune Landscape Phenotyping in Tumors by Muliplex Imaging

**DOI:** 10.1101/2021.10.22.464548

**Authors:** Shabaz Sultan, Mark A. J. Gorris, Evgenia Martynova, Lieke L. van der Woude, Franka Buytenhuijs, Sandra van Wilpe, Kiek Verrijp, Carl G. Figdor, I. Jolanda M. de Vries, Johannes Textor

## Abstract

Tissue specimens taken from primary tumors or metastases contain important information for diagnosis and treatment of cancer patients. Multiplex imaging allows *in situ* visualization of heterogeneous cell populations, such as immune cells, in tissue samples. Most image processing pipelines first segment cell boundaries and then measure marker expression to assign cell phenotypes. In dense tissue environments, this segmentation-first approach can be inaccurate due to segmentation errors or overlapping cells. Here we introduce the machine learning pipeline “ImmuNet” that identifies positions and phenotypes of cells without segmenting them. ImmuNet is easy to train: human annotators only need to click on an immune cell and score its expression of each marker. This approach allowed us to annotate 34,458 cells. We show that ImmuNet consistently outperforms a state-of-the-art segmentation-based pipeline for multiplex immunohistochemistry analysis across tissue types, cell types and tissue densities, achieving error rates below 5-10% on challenging detection and phenotyping tasks. We externally validate ImmuNet results by comparing them to flow cytometric measurements from the same tissue. In summary, ImmuNet is an effective, simpler alternative to segmentation-based approaches when only cell positions and phenotypes, but not their shapes, are required for downstream analyses. Thus, ImmuNet helps researchers to analyze multiplex tissue images more easily and accurately.

## Introduction

Tissue samples provide key information about the manifestation and progression of many diseases. In clinical oncology, histopathological examinations serve as an important basis for cancer diagnosis, treatment response monitoring, and relapse detection. There are also intensive ongoing efforts to develop histological biomarkers for selecting the appropriate treatment for cancer patients. Traditionally, tissue specimens are evaluated manually by trained pathologists, but machine learning (ML) systems are being developed for automating some aspects of tissue evaluation and improving the objectivity, reproducibility, and scalability of these aspects of histopathology.^1^

A core task of histopathological analysis is the localization of different types of cells. Many types of cells, such as epithelial cells and cancer cells, can be accurately identified based on morphological aspects like size, shape, or nuclear atypia. Leukocytes differ much less in morphology and need to be identified based on the expression of marker proteins. Such cells come in many flavors that require combinations of multiple markers for proper identification; for instance, T cells alone can be grouped in up to 10 major subsets, several of which can be sub-divided further.^2^ Accurate identification of immune cell subsets is especially important in the context of cancer immunotherapies, as different immune cell subtypes perform very different functions within the tumor microenvironment. Several multiplex imaging techniques such as multiplex immunohistochemistry (mIHC),^3^ co-detection by indexing (CODEX),^4^ cytometry by time of flight (CyTOF),^5^ or NanoString’s digital spatial profiling^6^ have been developed to allow *in situ* mapping of immune cell subsets. All these techniques deliver multi-channel images that typically consist of a nuclear stain (such as DAPI) together with nuclear, cytoplasm, or membrane markers to identify cell locations and phenotypes.

All current major analytic platforms are based on cell segmentation followed by quantifying the expression of each marker on the segmented cells.^7^ Indeed, a common view is that “it is impossible to extract statistics on cellular fluorescence or geometry with single-cell resolution”^8^ without an initial segmentation step. Variations of this approach subdivide each cell further into, for example, nucleus and membrane components; this allows to more accurately measure the expression of markers localized to that component of the cell. However, segmentation is a notoriously difficult problem in biomedical image processing, especially in dense tissue specimens.^9^ The biomedical imaging community has devoted significant efforts to cell segmentation, which has been the subject of several benchmark datasets and algorithm competitions (such as the weakly supervised cell segmentation challenge organized at the Neurips 2022 ML conference^10^). However, even near-perfect segmentation algorithms are affected by the fundamental issue that cells in dense tissues may overlap – which is unfortunately very often the case when it comes to immune cells in various tissues. Strategies to address this problem include dissolving the tissue^9^ – which loses important spatial information – and post-hoc corrections of expression profiles similar to compensation approaches in flow cytometry.^4^ In this paper, we take a fundamentally different approach: we develop a *segmentation-free* analysis pipeline^11^ that treats cell localization and phenotyping as one integrated problem and does not rely on segmentation as an initial step. We propose an ML pipeline that implements this approach and show that it achieves accurate results while being considerably easier to develop and train than segmentation-based pipelines.

## Results

### Segmentation-based phenotyping fails in dense tissues even when segmentation is perfect

Cell segmentation is a problem that has been studied for decades.^12^ Nowadays, end-to-end ML^8^ has largely replaced traditional approaches to cell segmentation such as the Watershed algorithm.^13^ In our experience, the StarDist ML pipeline for segmentation^14, 15^ often achieves excellent performance. Issues that complicate segmentation and subsequent phenotyping include: spectral unmixing effects, where channels are not separated well from each other; steric hindrance, where cells that are already stained with one antibody may become less efficient targets for subsequent antibodies;^3^ and high tissue density that leads to overlap between adjacent cells even in thinly cut slices. In particular, classic segmentation algorithms assign each pixel to one cell (or background) and are therefore not well suited for dense tissues.

To assess the need for developing a segmentation-free analysis pipeline instead of building on existing segmentation-based approaches, we analyzed multispectral images generated by a computer simulation model. Unlike real images, such *in silico* generated images have an available “ground truth”: we know exactly to which cell (or cells in case of overlap) each pixel in the image belongs. Therefore, we can use such images to reason about the hypothetical situation in which we have a *perfect* segmentation algorithm available, which helps us to put an upper bound on the performance that any such approach can achieve. Specifically, to mimic real fluorescent histopathological images as closely as possible, we used the Cellular Potts modeling framework.^16–18^ We placed cells of realistic size (about 5-10µm in diameter) into a 3D space representing an unlabelled background structure, and “label” membrane, nucleus, or both, depending on the simulated cell type. We cut out 4µm thick slices from these simulations for downstream analysis (**Figure 1A**). We then simulated noisy expression of nuclear, cytoplasmic, and membrane markers on these cells and integrated the expression values along the Z axis to obtain simulated 2D multispectral images, which indeed had a striking similarity to real multispectral images (**Figure 1A,B**).

**Figure 1:**
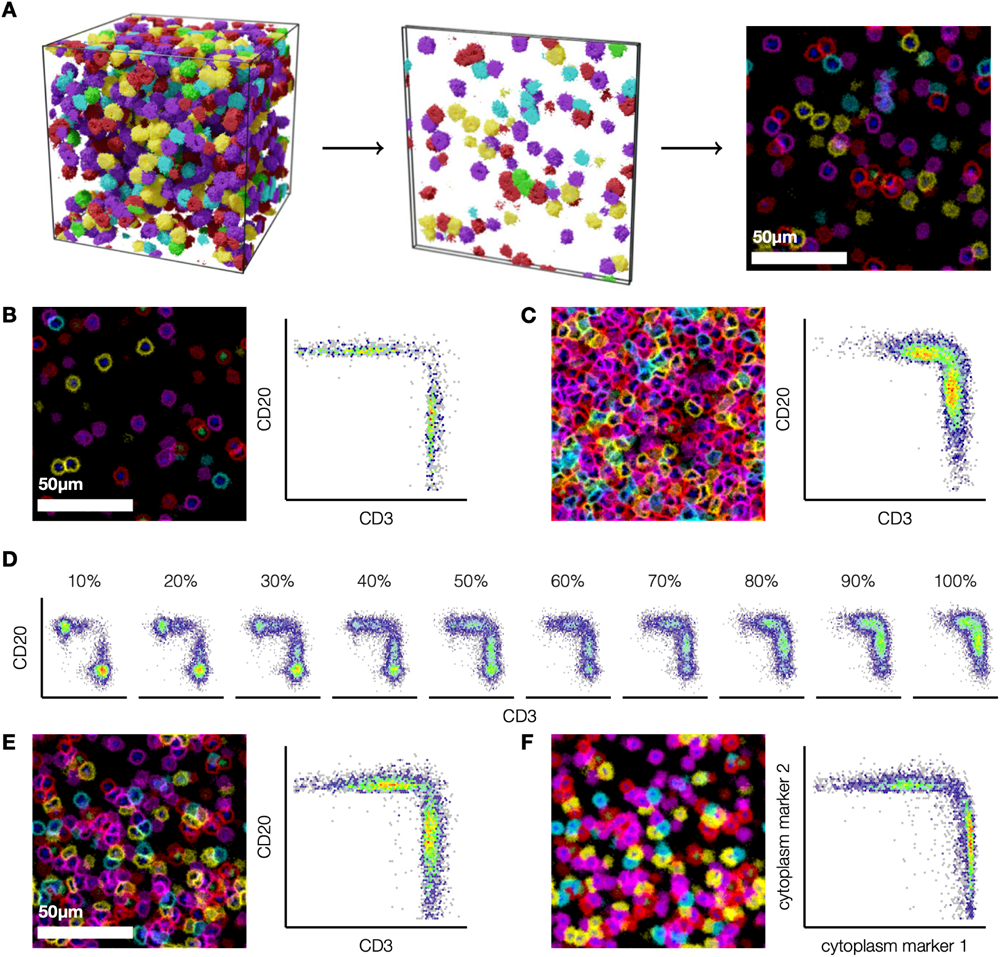
Segmentation-based phenotyping fails in dense simulated multiplex images. **(A)** To generate artificial immunohistochemistry images, we simulated cells at different densities within a 128_3_ µm3 volume, and cut 4µm thick *in silico* slices spaced 8µm apart. **(B,C)** Simulated tissue slices and corresponding scatterplots of CD3 and CD20 expression (membrane) on perfectly segmented cells at low (B) and high (C) cell densities. **(D)** At very high densities, individual cell populations are no longer identifiable (10% density: *∼*3,000 cells/mm2; 100% density: *∼*30,000 cells/mm2). **(E,F)** Compared to membrane-expressed markers (E), markers expressed in the entire cytoplasm (F) are less affected by noise and spillover from adjacent cells.

We used our simulated tissue images to assess the performance of segmentation-based phenotyping when using membrane markers. To this end, we generate simple flow-cytometry-like scatterplots of marker expression measured on segmented cells at varying densities, where we considered two membrane markers to represent CD3 and CD20 expression (**Figure 1B,C**). As expected, this approach worked very well at low and medium cell densities (<80%; **Figure 1D**): the plot showed clearly separate cell populations, which would be easy to classify in downstream analysis. However, at densities where cells overlapped, the separation between the different populations on the plots disappeared, creating the appearance of a single population with a continuum of expression of both markers. While it would still be possible to place arbitrary thresholds on these expression values to extract subpopulations, this approach would now either risk ignoring a substantial proportion of the cells, or misclassifying cells in the “double-positive” area. The problem was alleviated but not eliminated when we considered cytoplasm-based markers, which are less affected by cell overlap (**Figure 1E,F**).

In summary, our simulated data demonstrate that even if a perfect cell segmentation algorithm was available, segmentation-based phenotyping would still be difficult in dense tissues where expression readouts, especially of membrane markers, spill over between adjacent cells. This observation motivated us to develop a segmentation-free approach that treats cell phenotyping as a first-class problem rather than a downstream step of cell segmentation.

### A machine learning pipeline for direct phenotyping without segmentation

We develop our ML pipeline based on multiplex immunohistochemistry (mIHC) data of formalin-fixed paraffin-embedded (FFPE) tissue. Specifically, we employed mIHC using the Opal tyramide signal amplification technique and multispectral imaging.^19^ When used in conjunction with the Vectra 3 system or its successor, the Polaris, this method can combine 6 markers or more within one FFPE tissue section. However, because of the serial staining protocol, panels have to be optimized carefully.^3^ Using this technique, we developed a seven-color lymphocyte panel to detect different lymphocyte populations within tissue consisting of CD3, FOXP3, CD8, CD45RO, CD20, a tumor marker (such as pan-cytokeratin or a melanoma-specific antibody cocktail), and DAPI (**Figure 2A**).

**Figure 2:**
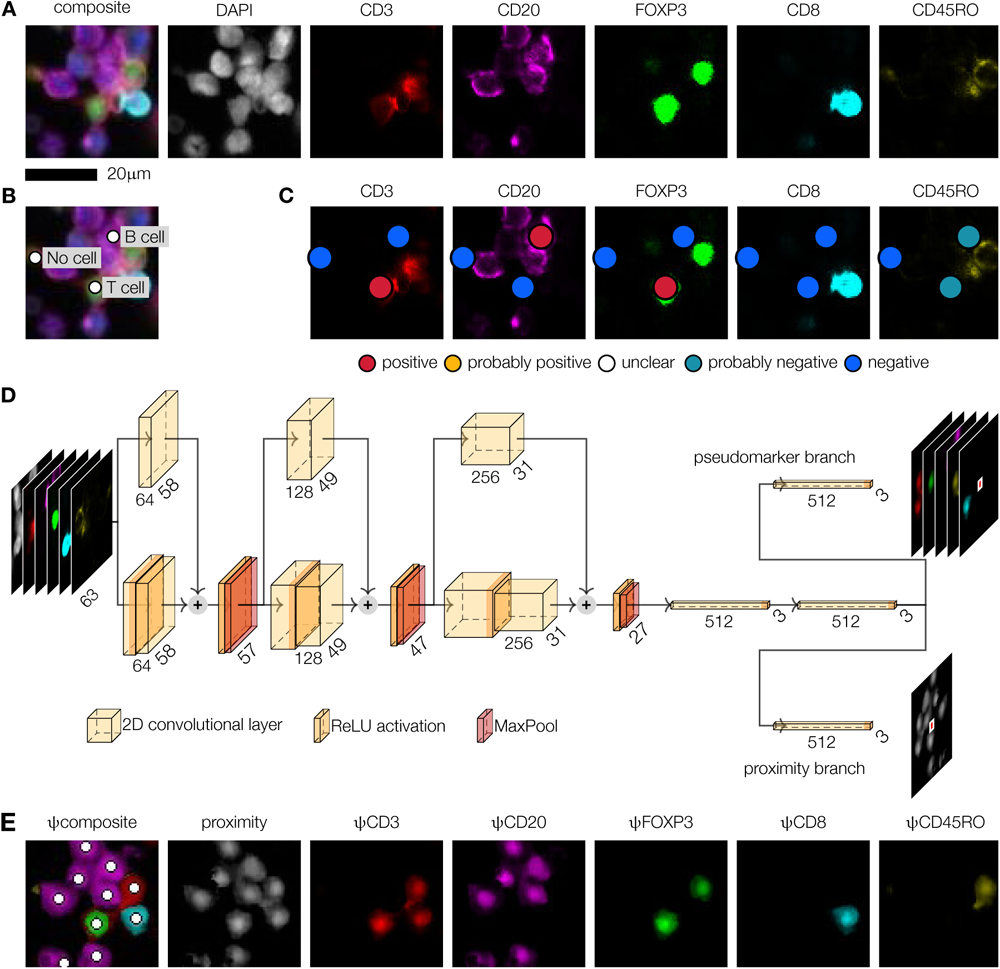
ImmuNet, an artificial neural network architecture for segmentation-free detection and phenotyping. **(A)** Multiplex immunohistochemistry imaging using a DAPI nuclear stain and a 5-marker panel designed to identify cytotoxic (CD8), regulatory (FOXP3) and memory (CD45RO) T cells (CD3) as well as B cells (CD20). **(B)** “Click” annotations of the locations of two cells and one background annotation (no cell). **(C)** “Decorations” of the annotations shown in (B) specifying the annotator’s certainty that each marker is expressed or not on the corresponding cell. **(D)** ImmuNet architecture consisting of 9 convolutional layers arranged in 3 blocks with skip connections, followed by a convolutional layer to reduce feature map size to 3×3, a fully connected layer and 2 output branches of a fully connected layer each. The network is trained on 63×63×7 input images and generates 3×3 output matrices containing the distance to the nearest cell (proximity branch) and the expression of each marker on the nearest cell (pseudomarker branch). **(E)** Output of the ImmuNet network on the input shown in (A). White circles show cell positions detected by Laplacian of Gaussian post-processing of the proximity map.

In previous studies, we used the software inForm (PerkinElmer) in conjunction with in-house developed down-stream quality control and analysis software to segment and phenotype cells in mIHC images.^3, 20–25^ Given our familiarity with this pipeline and our experience in fine-tuning it to specific tissues, we use it throughout this paper as a baseline method for comparison. The inForm software uses an ML algorithm to assign every pixel in the image to at most one cell, and subdivides each cell into compartments such as “nucleus” and “membrane”. Subsequently, it extracts marker expression information for each channel (e.g., mean expression, range of expression, variance of expression) along with morphological features such as size and shape indices. Users can manually annotate cells with known phenotypes to train a multinomial logistic regression classifier model that assigns a phenotype to each cell.^26^ As we will show, this approach works quite well on average, as long as the software parameters are appropriately tuned. However, as we will also demonstrate, the performance can degrade substantially for some cell types.

As is common in ML, we started by formulating our task in terms of the desired input and output. Existing neural network architectures for cell segmentation are typically based on images containing manually drawn cell outlines. Generating such cell boundary annotations is a time-consuming task. While sparse tissues are easier to annotate, networks trained on such images may perform poorly on dense structures they have not seen during training. For these reasons, we formulated two key requirements for our design: (1) users should only have to annotate the location of each cell (click annotation) instead of its entire shape (polygon annotation), given that we do not intend to use the shapes anyway; (2) users should not have to annotate training images fully, because even for a human expert it is difficult to identify every cell in dense tissues. We developed a custom annotation tool that allows users to place annotations simply by clicking on the center of cells of interest (**Figure 2B**). In a second step we call “decoration”, users can verify and finetune the locations of the annotated cells and rank the expression of each phenotyping marker on a five-point Likert scale (**Figure 2C**). Importantly, the Likert scale represents the user’s certainty that a cell expresses or does not express a certain marker rather than a qualitative judgment on expression intensity. This allows annotators to specify that they are uncertain about some cases, which can then be resolved by discussing these cases in a larger team of annotators and getting input from experts. In our case, such discussions took place only for a small amount (tens) of cases.

We then designed an artificial neural network (ANN) architecture that processes the location and phenotype annotations to generate two types of output per pixel: (1) the proximity of this pixel to the nearest center of a cell; (2) the expression of each phenotyping marker on the nearest cell (**Figure 2D,E**, **Supplementary Table 2)**. The network has a fully convolutional structure (**Figure 2D**, **Supplementary Table 2**) that allows it to generate whole-image predictions during inference despite generating only small patches of output during training (we use a patch size of 3×3 pixels to encode at least some information on the smoothness of the proximity function). This setup makes it straightforward to process sparsely annotated data: only pixels in close proximity to annotated cells are considered during training. Earlier prototype versions of our ANN architecture were inspired by the DeepCell network,^8^ although the final architecture turned out very different. We also incorporated a key idea of Wang et. al.,^27^ who trained a network on a similar distance transformation of cell locations.

Hence, our ANN architecture, which we called ImmuNet, generates maps that encode information about cell location and phenotypes, but not cell shape. These maps can be processed using any object detection algorithm. We found a simple Laplacian of Gaussian (LoG) blob detection algorithm to work well for our purposes (**Figure 2E**). The final output is a list of spatial coordinates of each detected cell and its expression of each marker quantified by what we call “pseudomarkers” (ψ). This kind of data is familiar to many biologists as it closely resembles the output of flow cytometers, but with added spatial information. Indeed, we found that converting ImmuNet data to the flow cytometry standard (FCS) format was an effective way to allow users to explore their multi-dimensional mIHC data using familiar software.

### Training and evaluation of ImmuNet on different types of tissues

Having defined our network architecture, we proceeded to collect data for annotation, training, hyperparameter tuning, and evaluation. To these ends, we created a database consisting of whole-slide mIHC images from four different types of human tumor samples (bladder cancer, lung cancer, melanoma, and prostate cancer; see Methods) as well as tonsil material from tonsillectomies (**Figure 3A**). All samples were stained using our T cell panel mentioned above except for the prostate samples, where we did not use the CD20 channel.

**Figure 3:**
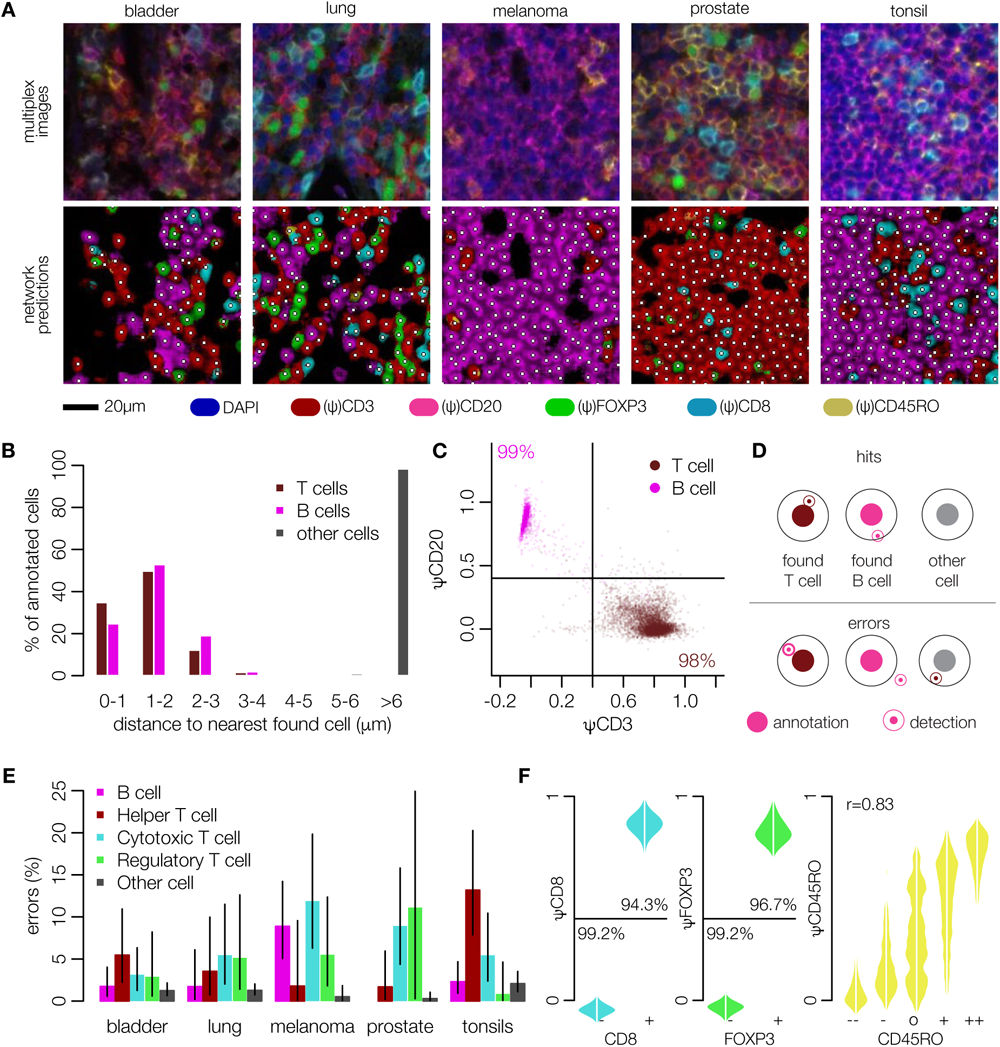
Identifying and phenotyping B-and T cells using ImmuNet. **(A)** Representative input images from 4 different types of tumor samples and a tonsil sample, focusing on dense lymphocyte clusters. **(B)** Distribution of distances between annotated cells and cells identified by ImmuNet. **(C)** Expression of the CD3 and CD20 pseudomarkers (ψ) on detected cells. **(D)** Definition of correct and faulty detections used in **(E)**, which shows the error rates of ImmuNet per tissue type compared to our baseline method. The detection radius used is 3.5µm. Error bars: 95% confidence intervals (Clopper-Pearson method). The wide confidence interval for regulatory T cells in prostate is a result of the low number of such cells found in prostate samples. **(F)** Distribution of pseudomarkers on detected cells compared to annotated marker expression on the nearest annotated cell. For CD8 and FOXP3, we grouped weakly and strongly positive or negative annotations.

Using our custom-built browser-based tools, we annotated and decorated thousands of lymphocytes of various types. In addition, we used “background” annotations, which include tumor cells, other cells – those which do not express any of the lymphocyte markers – and sites without a cell to provide the model with negative examples. To enrich our set of annotations for difficult cases, we performed several rounds of model training. At the end of each round, the output of the network was visualized and manually inspected to find areas where the network was making mistakes. Then, additional annotations were made at the problematic sites and the model was retrained. We stopped when we had accumulated 34,458 cell annotations, at which point we no longer found obvious problems with our network by visual inspection. We also trained baseline inForm segmentation and phenotyping algorithms on the same data, which unlike the ImmuNet approach required training a dedicated algorithm for each tissue type, and sometimes multiple algorithms per tissue type if there were substantial differences between batches (such as changed microscope configuration settings).

Since our multispectral images consisted of stitched individual tiles (1332×996 pixels), we separated the annotated tiles into training and validation sets to evaluate generalization to unseen tiles. Specifically, we used 20% of the annotations for validation and balanced the distribution of tissue types and phenotypes between training and validation data. The final training set comprised 27,888 annotations (lymphocytes, tumor cells and other cells), leaving 6,570 annotations for our initial validation; additional validation steps were also performed, and are described later in this paper. The training data were used to tune LoG parameters and to define hyperparameters as follows: after training our network and setting the LoG parameters, we measured the distance to the nearest detected cell for each annotated cell (**Figure 3B**). For the vast majority of annotated T-and B cells, ImmuNet detected a lymphocyte no further than 3µm away, which was rarely the case for non-lymphocyte annotations such as tumor cells or stromal cells. The expression of the CD3 and CD20 pseudomarkers corresponded closely to the annotated phenotype: for 98% of the annotated T cells and 99% of the B cells, the closest detected cell expressed the corresponding pseudo-marker – but not the other pseudomarker – at an intensity of 0.4 or higher (**Figure 3C**). Thus, in the training data, most of the annotated cells were correctly detected by the ImmuNet pipeline when using a distance cutoff of µm and an intensity cutoff of 0.4.

We next determined the pipeline’s performance in the detection of T cell phenotypes in the validation data. While making annotations, we generally found it easy to decide on positivity for CD20, CD3, CD8 and FOXP3 markers. We therefore binarized the Likert scale annotations to “positive” and “negative” for these markers and discarded the few annotations for which this distinction could not be made. The status of the CD45RO marker was more difficult to assess, so we kept the original Likert values. Likewise, we used the established cutoff of 0.4 to binarize the predicted pseudomarker expression values except for ψCD45RO. In this way, we defined five different phenotypes as a basis for evaluation: B cells (CD20^+^CD3^-^CD8^-^FOXP3^-^), helper T cells (“Th” for short; CD3^+^CD8^-^FOXP3^-^CD20^-^), cytotoxic T lymphocytes (“CTL”; CD3^+^CD8^+^FOXP3^-^CD20^-^), regulatory T cells (“Treg”; CD3^+^FOXP3^+^CD8^-^CD20^-^), and other cells (CD20^-^CD3^-^CD8^-^FOXP3^-^). We treated any other combination of predicted CD20, CD3, CD8, and FOXP3 levels as an invalid prediction. We then evaluated the entire cell detection and phenotyping process as follows: for each annotated B-or T cell, we require the closest detected lymphocyte to be no further than 3.5µm away, and it must have the same phenotype. For other annotated cells, no B-or T cell must be detected by the network within a 3.5µm radius (**Figure 3D**).

Using these definitions, we found that the ImmuNet error rate in the validation data was below 10% for B cells and for most T cell subtypes (21 out of 24 categories tested; **Figure 3E**), and below 5% for 14 out of the 24 categories tested. False positives (i.e., “other cell” annotations where the network predicted a lymphocyte in close proximity) were also acceptably low with tonsils showing the largest false positive rate at only 2.1%. For every possible combination of marker and tissue, comparing these values to the error rates of our baseline inForm algorithms (which were also trained on cell annotations collected from these datasets) showed that ImmuNet significantly outperforms inForm in lymphocytes detection and phenotyping: error rates measured on the same validation dataset were above 20% for B cells and above 10% for T cell subtypes across all tissue types, sometimes reaching values as high as 40% even after re-training each inForm algorithm twice to optimize performance (Methods; **Supplementary Figure 1)**.

Analyzing the pseudomarker values in more detail for T cell subsets likewise showed excellent agreement between annotated and predicted expression (**Figure 3F**, left). Since the annotations of the CD45RO marker were more uncertain, we visualized pseudomarker distributions for each value of the Likert scale (**Figure 3F**, right), which confirmed that it would not be appropriate to use a binary cutoff for this marker, as too much information would be lost. Instead, we characterized the agreement between annotations and predictions using the Pearson correlation, which was reasonably high (0.83). Importantly, when an annotator was more certain (first and last level of the Likert scale), the pipeline’s prediction matched the annotator’s judgement in most cases.

Taken together, these results showed that ImmuNet can detect and phenotype B cells, helper T cells, cytotoxic T cells and regulatory T cells using a pseudomarker expression cutoff of 0.4, whereas CD45RO expression needs to be evaluated on a continuum, avoiding binarization. Error rates were consistently below 10% and often below 5%, a major improvement compared to our previously used baseline method.

### Validation on fully annotated regions of interest

Encouraged by the low error rates found in this initial analysis, we performed a second round of testing designed to better detect additional potential issues. First, even low false positive rates might still be problematic if entire slides are analyzed that contain almost no lymphocytes: in such cases, the majority of detected lymphocytes could be errors. Second, we noticed that hypersegmentation of cells – where a single cell is being detected as multiple cells – was a common problem with our inForm algorithms and was not being picked up in our initial test due to our focus on single cells. Additionally, when making sparse annotations, annotators might be biased towards selection of easier cases. Therefore, we next fully annotated 229 of regions of interest (ROIs; **Figure 4A**) representing different tissues, cell compositions, and density levels.

**Figure 4:**
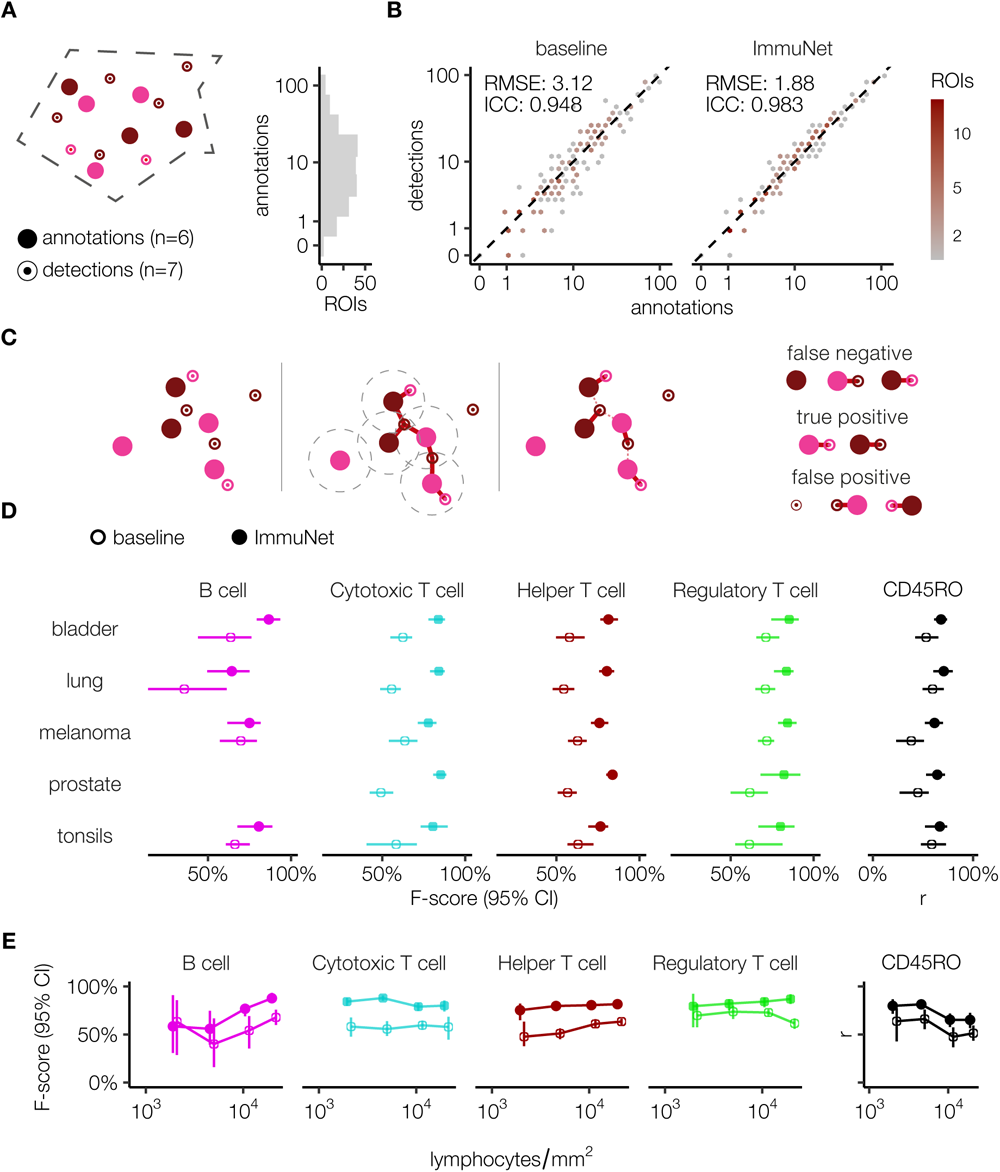
Validation of ImmuNet lymphocyte detection across tissue environments, phenotypes, and densities. **(A)** Sketch of an ROI with human annotations and detections made by ImmuNet, with different colors representing different phenotypes; the histogram shows the distribution of the number of annotations per ROI. **(B)** Number of annotations and detections for baseline algorithm and ImmuNet, visualized using hexagonal binning. **(C)** Illustration of the definition of TPs, FPs and FNs by optimal pairwise matching of annotations to detections. **(D)** ImmuNet and inForm performance across tissue types and cell phenotypes. Error bars: bootstrapped 95% CIs (2000 replicates). **(E)** ImmuNet and inForm performance across density profiles and cell phenotypes. Error bars: bootstrapped 95% CIs (2000 replicates).

First, we determined the agreement between the number of annotated and detected cells in each ROI by calculating the intraclass correlation coefficient (ICC) – a measure of interrater agreement that ranges from -1 (perfect disagreement) to 1 (perfect agreement) – and root mean square error (RMSE) between these values for ImmuNet and the inForm baseline (**Figure 4B**). Whereas inForm achieved an ICC of 0.948 and a RMSE of 3.12, ImmuNet improved both metrics, increasing the ICC to 0.983 and reducing the RMSE to 1.88 cells. **Figure 4B** highlights that inForm tends to miss lymphocytes frequently, and that this may on average be a bigger issue than hypersegmentation. By contrast, ImmuNet seems to over-and underestimate the number of lymphocytes similarly often.

Next, we assessed the accuracy of cell phenotyping and localization inside ROIs. For B cells, Helper T cells, Cytotoxic T cells and Regulatory T cells, we used F-scores as an evaluation metric, whereas the accuracy of the predicted CD45RO expression was evaluated using the Pearson correlation of Likert scale annotation with predicted positivity (inForm instead performed a binary CD45RO classification). To compute these metrics we needed to match detections and annotations in each ROI to each other. For this, we used a standard maximum matching algorithm,^28^ which matches as many cells as possible while respecting the distance cutoff of 3.5µm. This matching allows us to calculate the number of true positives (TPs), false positives (FPs) and false negatives (FNs) in each ROI as illustrated in **Figure 4C**: We consider the predicted cell to be a TP if it is matched with an annotation of the same phenotype. An FP is defined as a detection that is unmatched or matched to an annotation with a different phenotype. Conversely, unmatched annotations and annotations matched with a prediction of a different phenotype are considered TNs. For the CD45RO marker, the Pearson correlation between the Likert annotation and the pseudomarker expression was computed for the matched CD3^+^ annotations and detections.

We performed this evaluation across different tissue types (**Figure 4D**) and density profiles (**Figure 4E**). Density profiles were obtained by binning the log-scaled range of annotated lymphocyte densities in all ROIs. F-scores for phenotypes and tissue environments (density profiles) are calculated by bootstrapping upon aggregating TPs, FPs and FNs across all ROIs. Pearson correlation for the CD45RO marker is calculated in the same way. These results demonstrate that ImmuNet outperforms inForm across lymphocytes phenotypes, different tissue types and density profiles. Importantly, ImmuNet’s performance did not drop much at higher tissue densities, except for prediction of CD45RO expression. By contrast, we did notice an interesting drop in inForm’s performance for T regulatory cells at high densities. Since FOXP3 is a nuclear marker, its measurements should still be fairly reliable even at high densities. However, it does rely on a reasonable initial segmentation, since regulatory T cells are rare and often surrounded by cells of other types.

Taken together, our analyses on two differently designed validation datasets demonstrated robust performance of ImmuNet across tissue types and cell densities and significantly outperformed the baseline “inForm”, our previously preferred method.

### External validation of ImmuNet results using flow cytometry measurements of the same tissue

Up to this point, our performance evaluations were based on manually annotated cells. While this is a standard procedure in ML, we noticed that human annotators did not always agree in their judgements about cell counts and phenotypes; upon closer examination of prediction errors, we repeatedly found cases where the ground truth was in fact debatable. We therefore designed an experiment to compare our IHC phenotyping data to external reference measurements rather than human annotations. Flow cytometry is a mature and widely used non-spatial method for cell phenotyping. Because cells in a flow cytometer are dissociated and imaged one by one (rare duplicates can be filtered out in post-processing), and the entire outside of a cell is accessible to the cytometer, marker expression can be measured more reliably compared to mIHC imaging. We therefore decided to use flow cytometry as an external control for the relative lymphocyte phenotype abundances estimated by mIHC-based phenotyping. To this end, we obtained fresh human tonsil tissue. Tonsils are a useful test case for our analysis because they contain extremely densely packed B-and T cell areas that are notoriously difficult to process for segmentation algorithms. For further processing, we split each tonsil in half (**Figure 5A**). One half of each tonsil was dissociated into single cells and analyzed by both flow cytometry and an FFPE AgarCyto cell block preparation^29^ subjected to mIHC. The other half was directly FFPE and subjected to mIHC. We then quantified the number of B cells, T helper cells, cytotoxic T cells and regulatory T cells as a proportion of all B-and T cells for each of the three preparations.

**Figure 5:**
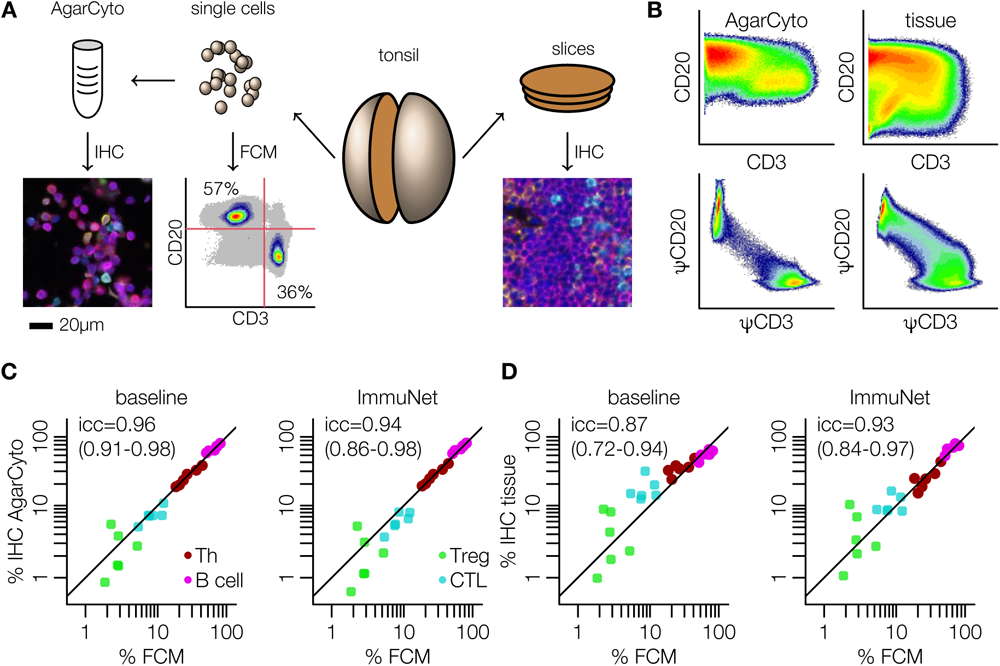
External validation of ImmuNet-derived phenotype abundances. **(A)** Tonsils were cut in half with one half sliced and imaged by IHC, and the other half dissolved and further processed by flow cytometry or an AgarCyto preparation, which was also imaged by IHC. **(B)** Directly measured expression of CD3 and CD20 on segmented cells from the IHC images compared to the expression of the corresponding pseudomarkers on ImmuNet-detected cells. **(C,D)** Cells in AgarCyto (C) or direct FFPE (D) mIHC images were either phenotyped using the multiparametric classifier implemented in inForm (baseline), or using a threshold of 0.4 on the ImmuNet pseudomarkers. Concordance to flow cytometry measurements from the same tonsils is shown and quantified using the ICC and its 95% CI.

Two-dimensional plots of CD3 versus CD20 expression clearly showed distinct peaks representing T-and B cells for the flow cytometry data (**Figure 5A**). While two separate peaks were still somewhat apparent from the AgarCyto preparation analyzed with the inForm baseline method, these disappeared when directly imaging the dense tonsil tissue (**Figure 5B**), resembling our initial findings on simulated data (**Figure 1B**). By contrast, separate peaks in the expression of ImmuNet pseudomarkers were still readily identifiable for both the AgarCyto preparation and direct FFPE tissue imaging (**Figure 5B**). To phenotype the cells based on their expression profiles, we again used the positivity threshold of 0.4 identified previously for CD3, CD8, FOXP3 and CD20 (**Figure 3**). Given that a similar direct thresholding of expression markers would not seem sensible for the inForm data (**Figure 5B**), we instead trained and applied the inForm phenotyping classifier, which can take many additional features into account, to determine the baseline performance.

Reassuringly, our analysis showed good agreement between flow cytometry and AgarCyto-mIHC measurements in both the baseline analysis and in the ImmuNet data (**Figure 5C**). On the directly FFPE mIHC images, the agreement of the baseline method with flow cytometry data degraded somewhat from ICC=0.96 to ICC=0.87, mainly due to an apparent overcounting of CTLs, while the ImmuNet-based agreement remained very similar (ICC=0.94 versus ICC=0.93). We noticed that these agreement measurements were strongly influenced by the Treg population, which was hard to gate and therefore less reliable on the flow cytometry data (**Supplementary Figure 2A**). We therefore performed the same analysis without the Treg population, which gave similar results for ImmuNet but a more marked loss in performance for the baseline method on the tissue images (**Supplementary Figure 2B**). An important caveat of this analysis is that one does not necessarily expect perfect agreement between flow cytometry and tissue images because suspension could lead to the partial loss of some cell populations. In line with this, the agreement between the different analysis methods (baseline and ImmuNet) on the same imaging data was generally higher than agreement between flow cytometry and tissue data (**Supplementary Figure 2C**).

In summary, our comparison of mIHC phenotyping compared to flow cytometry analysis of the same tissue showed generally good agreement between the two methods, provided that cells were easy to “gate” in both methods. Our analysis suggested that immune cell phenotyping achieved by ImmuNet was at least on par that achieved by our segmentation-based baseline method, which appeared to struggle with oversegmentation of CTLs in the dense tonsil tissue.

### Spatial analysis of tumor microenvironments reveals distinct infiltration patterns and local interactions of lymphocytes

To illustrate the applicability of ImmuNet approach in cancer research, we performed a spatially resolved analysis of tumor-infiltrating lymphocytes (TILs) in the tumor microenvironment. Such analyses are common in the literature because TIL density has prognostic or predictive value in different types of cancer including melanoma,^30, 31^ lung cancer,^32^ and colorectal cancer.^33^ Based on TIL density within and around the tumor, tissue samples often described as “hot” (TIL-rich), “excluded” (TIL-rich in stroma, but not tumor), and “ignored” (TIL-poor),^34^ or TIL distribution is examined using more refined measures of effective infiltration.^20, 35^ Multiplex imaging techniques add phenotype information to such approaches, allowing the analysis of interactions between different types of cells.^36^

We used ImmuNet to analyze TILs in our whole-slide mIHC images of bladder cancer, lung cancer, melanoma, and prostate cancer tissue specimens (**Figure 6A**), applying a simple threshold-based segmentation algorithm to distinguish tumor and stroma (Methods; **Supplementary Figure 3**). Similar to a previous study,^35^ we measured the density of T cells (combining all phenotypes) and B cells in the invasive margin, defined as a region within 100µm around the tumor-stroma boundary. For bladder cancer, lung cancer, and melanoma, the density of TILs in stroma was consistently higher than or comparable to their density in tumor; these samples could all be described as “hot”, “excluded” or “ignored” based on the absolute density (**Figure 6B**). In the prostate cancer samples, there was a different picture where the density of cells outside the tumor was consistently lower.

**Figure 6:**
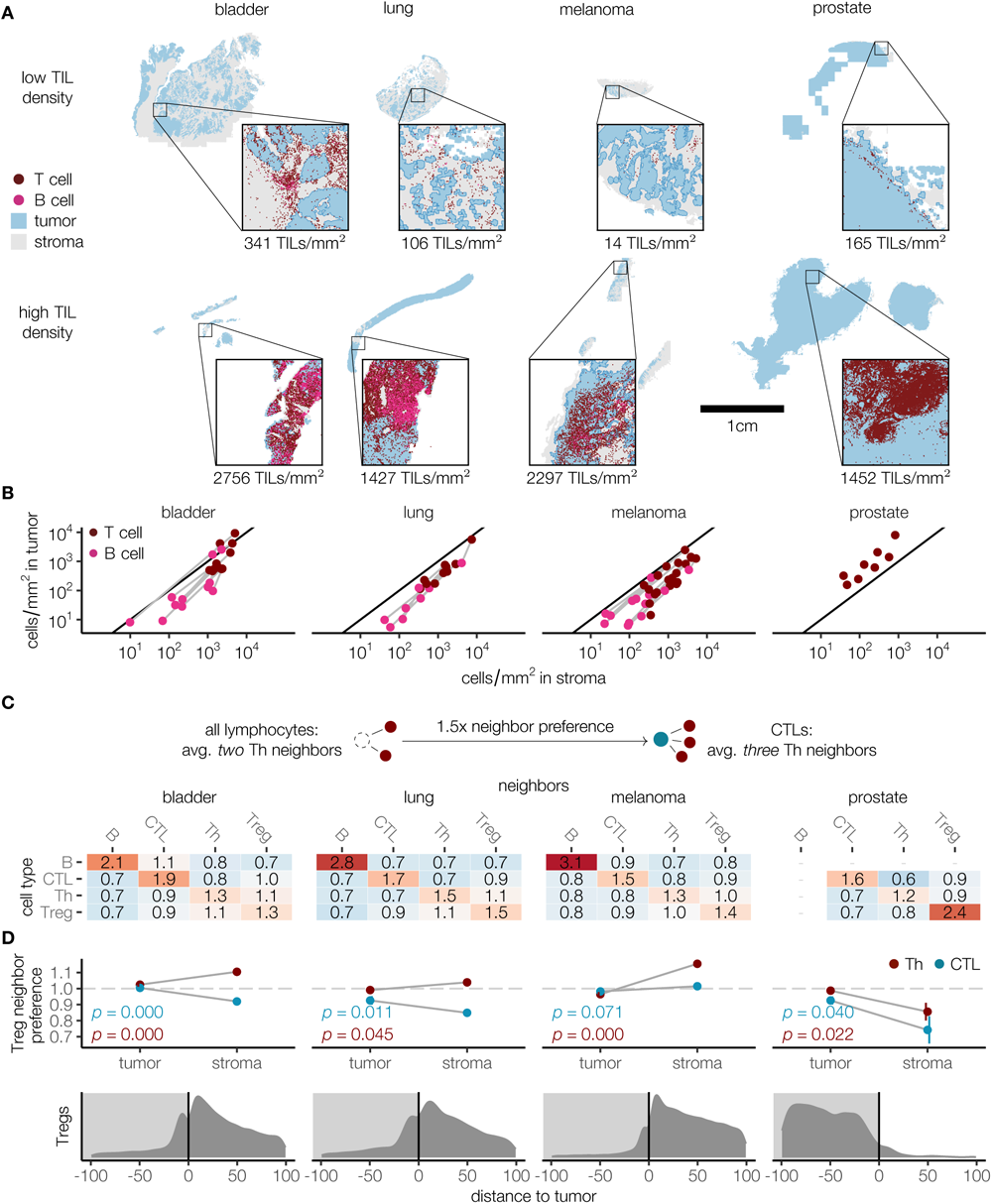
Quantifying immune cell infiltration and cell-cell interactions in the tumor microenvironment. **(A)** Representative sections in different tissue environments with low and high TIL density. **(B)** Density of B and T cells inside or outside the tumor (100μm from the border); prostate sections were not stained with CD20. **(C)** Analysis of neighboring cells, at up to 14μm between cell centers. Average occurrence of different cell types as neighbor is calculated for neighbors of lymphocytes in general, and for neighbors of specific cell types. Ratio between these numbers is deemed as preference of a certain type of cell to have another cell type as neighbor. **(D)** Stratified analysis of Treg neighbor preference by tissue type (top) and frequency of Tregs by distance to stroma/tumor border (negative numbers are inside the tumor). P-values from a T test of the null hypothesis of no difference between tissue type are shown. Bladder: N=11 whole-slide images; lung: N=10; melanoma: N=23; prostate: N=8.

Next, we assessed local cell-cell interactions between different lymphocytes on a smaller spatial scale. We designed a “neighbor preference” metric to quantify which cell types preferentially appear next to each other. The metric is calculated by determining how often a certain type of cell is present as neighbor of other lymphocytes in general (e.g., helper T cells as neighbors of all lymphocytes) and how often they are neighbor to a specific type of cell (e.g., helper T cells as neighbors to cytotoxic T cells specifically; **Figure 6C**). The ratio between these counts indicates how frequently two cells of the investigated types appear next to each other compared to a random assignment of cell types. This analysis revealed similar interaction patterns across the different tissue types (**Figure 6**): most cell types were preferentially close to cells of their own type (neighbor preference >1) whereas co-localization between cells of different types was less common (neighbor preference <1). B cells had overall the greatest tendency to co-localize with their own kind; this could be due to the presence of follicle-like structures in the tumor microenvironment.

As the regulatory T cells showed the weakest overall preference for or against other T cell phenotypes (as seen from the Th-Treg and Th-CTL values being close to 1), we stratified this interaction further by distinguishing between intra-and peritumoral cells (**Figure 6D**, top row). In all tissues, intratumoral regulatory T cells had no strong preference for helper T cells or cytotoxic T cells. In the stroma, however, they preferred helper T cell neighbors over cytotoxic T cells. For prostate cancer, we observed an apparent preference for homogeneous contacts, but this is likely a consequence of the very low number of peritumoral regulatory T cells in these samples (**Figure 6D**, bottom row).

In summary, the ImmuNet pipeline allowed us to quantify infiltration patterns in different environments, determine cell-cell interactions, and to stratify cell-cell interaction patterns by spatial compartment. This illustrates that cell segmentation is not always required to conduct spatial analyses of immune landscapes in tissues.

## Discussion

We have developed, implemented, trained, and tested ImmuNet, an ML pipeline for segmentation-free localization and phenotyping of immune cells in mIHC imaging. Although relatively little information is used to train ImmuNet compared to segmentation-based ML pipelines such as StarDist,^14^ we found it to perform well across diverse types of tissues, including very challenging dense environments. By design, ImmuNet is particularly well suited for applications where the cell shapes are not required for downstream analysis. This should often be the case for immune cells, because lymphocytes anyway tend to lose their physiological shape in dead tissue and round up. Indeed, our example analyses illustrate that sophisticated spatial information can still be extracted without cell shapes. However, for applications where cell shape is important, one could combine ImmuNet with a cell segmentation of the same tissue: the pseudo-markers could simply be added as additional channels to the image for further analysis in segmentation software such as CellProfiler^37^ or ilastik.^38^ Alternatively, a cell segmentation network like StarDist^14^ could be extended with additional branches to generate ImmuNet pseudomarkers alongside segmentation maps.

We built the ImmuNet pipeline shown in this paper specifically for our T cell-focused antibody panel; we have in the meantime applied it successfully to other panels including a dendritic cell panel (van der Hoorn *et al.*, in preparation). In practice, panels are adapted often, depending on the specific research question. A cautious approach would be to train a new ImmuNet from scratch for each panel. This is laborious but feasible given the relative ease at which large numbers of annotations can be collected – using our internal tooling, one person can typically annotate a few hundred cells per hour. However, when only one or two markers change, it may be effective to pool the data, especially if the alternative surface markers are of the same type (e.g., if one membrane marker is exchanged for another). Similarly, when some markers turn out to be unreliable in certain samples because of unspecific staining, one can still pool the data with other samples where the same marker is used but zero out the unreliable channel. In future research, we would like to investigate to what extent transfer learning strategies^39^ or image transformers^40^ could be employed to mix different channels in a more flexible manner. However, spillover between adjacent mIHC channels complicates developing such mix-and-match techniques; these could be easier to apply other multiplex imaging methods such as CyTOF, that are less affected by spillover effects. This may however not be straightforward for mIHC data given the spillover effects between adjacent channels frequently seen in such data and might be a more fruitful avenue for other multiplex imaging techniques that are less affected by spillover such as CyTOF.

In summary, ImmuNet is a simple but effective ML pipeline for cell detection and phenotyping in multiplex imaging; for example, we have already used ImmuNet to analyze melanoma,^41^ lung cancer,^42^ prostate cancer,^43^ bladder cancer,^44^ rectal cancer^45^ and other cohorts. Although we developed and tested ImmuNet for FFPE mIHC data, it should also be applicable to other multiplex imaging systems such as CyTOF, CODEX, and NanoString. We hope that ImmuNet pipelines will help researchers to generate more reliable phenotype maps of immune cells in tissue samples as a robust basis for exploratory, diagnostic, and prognostic applications of multiplex imaging technologies.

## Materials and Methods

### Ethics approval and consent to participate

The study of the melanoma material collected at the Radboudumc was officially deemed exempt from medical ethical approval by the local Radboudumc Medical Ethical Committee concurrent with Dutch legislation, as we used leftover coded material and patients are given the opportunity to object to their leftover material to be used in (clinical) research. Lung cancer samples were obtained from the PEMBRO-RT Phase 2 Trial^46^ at the Netherlands Cancer Institute, which was approved by the institutional review board or independent ethics committee of the Netherlands Cancer Institute–Antoni van Leeuwenhoek Hospital, Amsterdam. All lung cancer patients provided written informed consent and consented to further analysis of patient material collected prior to and during the PEMBRO-RT trial. The research on prostate and bladder cancer samples was approved by the local Radboudumc Medical Ethical Committee (file number 2017-3934). All patients provided written informed consent to scientific use of archival tissue, unless deceased.

### Simulation of artificial tissues

To generate synthetic *in silico* multiplex images, we used the cellular Potts model.^18^ Briefly, cells are randomly placed in a 128_3_ μm_3_ volume and simulated at a resolution of 0.5_3_ μm_3_ per voxel, matching the resolution of real multiplex images. Cells consist of two compartments, a nucleus and a surrounding cytoplasm region. Cells were randomly assigned a phenotype, with associated nuclear and membrane expressed markers matching real cells. The simulation operates by placing seed voxels of cytoplasm for each cell at a random location, and letting them circularize for 25 Monte Carlo steps. Then a nucleus seed voxel is placed inside each cell and the simulation is run for a further 50 Monte Carlo steps so cells can settle into their final shape. Settings controlling size of nucleus and cytoplasm per cell and adhesion strengths are described in Supplementary Table 1. Simulation temperature was set to *T* = 20.

To simulate membrane expressed markers, all voxels on the outer layer of a cell’s cytoplasm compartment (i.e., voxels having a neighbor that does not belong to the cell) are found and marked as membrane. All voxels inside a cell’s nucleus are used for nuclear expressed markers. An 8 voxel thick slice is taken from the simulation volume, corresponding to a 4µm tissue slice, matching our imaged tumor tissue slides. For each simulated marker, expression is simulated in either membrane or nuclear voxels, and signals are integrated along the viewing direction to construct an image. An exact cell segmentation mask is extracted from simulation data from the middle of the 8 voxel thick slice.

### Human material

Tonsils were collected from patients undergoing routine tonsillectomy at Canisius Wilhelmina Hospital in Nijmegen. Tonsils were stored at 4°C and processed within 24 hours. Tonsils were cut in halves of which one half was formalin fixed and paraffin embedded and the other half was processed into a single cell suspension. Fatty tissue was removed from the tonsil as much as possible with scalpels and placed into a gentleMACS C-tube (130-096-334, Miltenyi Biotec) with 5ml RPMI containing 0.3mg/ml Liberase (000000005401020001 Sigma) and 0.2mg/ml DNAse I (18068-015, Thermo Fisher). Tonsil tissue was roughly cut into smaller fragments using scissors and was further dissociated into a single cell suspension on the gentleMACS (130-096-334, Miltenyi Biotec) program “Multi_C_01_01” two times with a 15 minute incubation in between in a shaking water bath for 15 min-utes at 37°C. 1*×*10_6_ cells were used for flow cytometry measurements and 1.5*×*10_7_ cells were fixed and embedded in paraffin with the AgarCyto cell block preparation.^29^

Melanoma specimens from patients treated at the Radboud university medical center (Radboudumc) were randomly included based on the availability of a resection specimen. Lung cancer samples were collected at the Netherlands Cancer Institute in the PEMBRO-RT Phase 2 Randomized Clinical Trial.^46^ Bladder cancer samples were derived from patients who were treated for metastatic bladder cancer at the Radboudumc between 2016 and 2019. Archival tissue of both primary and metastatic tumor lesions was used. Prostate cancer samples are derived from patients that were treated in the Radboudumc and include archival tissue of both primary and metastatic tumor lesions.

### Flow cytometry

Single cells from tonsil (10_6_) were stained with Fixable Viability Dye eFluor™ 780 (eBioscience, 65-0865-18, 1:1000) for 20 minutes at 4°C. After wash steps, cells were incubated with a mix of anti-CD3-BV421 (BD Bioscience, 563798, clone SK7, 1:25), anti-CD8-PerCp (BD Bioscience, 345774, clone SK1, 1:5), anti-CD45RO-APC (BD Bioscience, 340438, clone UCHL-1, 1:25), and anti-CD20-PE (Biolegend, 302306, clone 2H7, 1:10) for 30 minutes at 4°C. Next cells were fixed, permeabilized with Foxp3/Transcription Factor Staining Buffer Set (eBioscience, 00-5523-00) and incubated with anti-FOXP3-alexa488 (eBioscience, 53-4776-42, clone PCH101, 1:8) 30 minutes at RT. Flow Cytometry was conducted with the FACS Verse (BD Biosciences). Flow cytometry data was analyzed using FlowJo software (v10, Tree Star).

### Multiplex immunohistochemistry staining

Sections of 4μm thickness (except for tonsil tissue, where we used 1μm) were cut from FFPE tissue blocks containing tonsil, melanoma and AgarCyto preparations respectively. The slides were subjected to sequential staining cycles as described before,^3^ although now automated using Opal 7-color Automation IHC Kit (NEL801001KT; PerkinElmer) on the BOND RX IHC & ISH Research Platform (Leica Biosystems) as described previously.^23^ All heat induced epitope retrievals were performed with Bond™ Epitope Retrieval 2 (AR9640, Leica Biosystems) for 20 minutes at 95°C. Blocking was performed with antibody diluent. Primary antibody incubations were performed for 1 hour at RT. All secondary antibody incubations were performed for 30 minutes at RT. mIHC was performed with anti-CD45RO (Thermo Scientific, MS-112, clone UCHL-1, 1:3000) and Opal620, anti-CD8 (Dako, M7103, clone C8/144B, 1:1600) and Opal690, anti-CD20 (ThermoFisher, MS-340, clone L26, 1:600) and Opal570, anti-CD3 (ThermoFisher, RM-9107, clone RM-9107, 1:400) and Opal520, FOXP3 (eBioscience Affymetrix, 14-4777, clone 236A/E7, 1:300) and Opal540. For prostate cancer samples, anti-CD56 (Cell Marque, 156R-94, clone MRQ-42, 1:500) was used with Opal570 instead of anti-CD20. However, for the purpose of this paper, we ignored the CD56 marker by setting the corresponding channel to 0 before processing.

To visualize tumor cells, melanoma tissues were stained in the end with a melanoma mix consisting of anti-HMB-45 (Cell Marque, 282M-9, clone HMB-45, 1:600), anti-Mart-1 (Cell Marque, 281M-8, clone A103, 1:300), anti-Tyrosinase (Cell Marque, 344M-9, clone T311, 1:200) and anti-SOX-10 (Cell Marque, 383R-1, clone EP268, 1:5000) and Opal650 to visualize tumor tissue. Tonsil, bladder and lung cancer tissues were finished with anti-pan cytokeratin (Abcam, ab86734, clone AE1/AE3 + 5D3, 1:1500) and Opal650 to visualize epithelial tissue. Finally, epithelial tissue in prostate cancer samples was visualized with a mix consisting of anti-pan cytokeratin (Abcam, ab86734, clone AE1/AE3 + 5D3, 1:1500), anti-EPCAM (Abcam, ab187372, clone VU-1D9, 1:1000) and anti-PSMA (Bio SB, BSB6349, clone EP192, 1:1000) and Opal650. Slides were counterstained with DAPI and mounted with Fluoromount-G (SouthernBiotech, 0100-01).

### Tissue imaging and data preparation

Slides were scanned using the PerkinElmer Vectra 3.0.4. Multispectral images were unmixed using spectral libraries build from images of single stained tissues for each reagent and unstained tissue using the inForm Advanced Image Analysis software (inForm 2.4.1, PerkinElmer). Per dataset, a selection of 1 to 2 representative original multispectral images per sample were used to train an inForm algorithm for tissue segmentation (into tumor/epithelium, stroma, background based on DAPI, tumor marker, and autofluorescence), cell segmentation, and phenotyping tool to recognize tumor cells, other cells, B cells and T cells populations (CD3^+^CD8^-^CD45RO^-^, CD3^+^CD8^-^CD45RO^+^, CD3^+^CD8^+^CD45RO^-^, CD3^+^CD8^+^CD45RO^+^, CD3^+^FOXP3^+^CD45RO^-^, CD3^+^FOXP3^+^CD45RO^+^). All settings applied to the training images were saved within an algorithm and batch analysis of whole slides was performed. After checking the performance of each batch analysis, erroneous images were added to the inForm algorithms and retrained to improve phenotyping of cells. A total of three iterations was performed per dataset to optimize the inForm algorithms. Component data files were used as input for ImmuNet.

### Artificial neural network

#### Rationale for network architecture choices

Analysis of cellular images has greatly advanced in recent years with the introduction of deep learning algorithms.^47^ Using these algorithms, cell segmentation can be formulated as a pixel-wise supervised learning task, where e.g. each pixel is classified as being part of a cell, a cell boundary or as background. Fully convolutional neural networks (FCNNs) are one method to perform pixel classification on whole images.^48^ For each pixel the FCNN gets to see a “receptive field” of surrounding pixels, allowing it to use local context to judge cell type and boundary shape per pixel. An important variant of this approach is U-Net.^49^ U-Net is designed for whole slide analysis; where FCNNs only see smaller structures in an image, U-net integrates both pixel level detail and large scale information to segment structures bigger than an FCNN can perceive.

We had initially compared FCNN and U-net architectures, and in early testing they performed comparably. We found the FCNN architecture a more natural fit for our sparse annotations, and hypothesized that because we segment cells that are small enough for an FCNN to observe, our task might not benefit greatly from U-Net’s ability to take larger environments into account. For these reasons, we decided to use an FCNN architecture, but we are planning to explore how this compares to U-net on our current data in future research.

In the early stages of our development process, we had considered and compared both location-based and cell shape-based annotations, taking the idea for location annotations from earlier studies on IHC data that used fully annotated training images.^50, 51^ During our early testing we found location-based annotations combined with a distance transformation to perform well, and therefore did not proceed with the much harder to obtain shape-based annotations. We experimented with splitting up distance and phenotype predictions in separate FCNNs, or have dedicated networks for B and T cells; however, in the end we found that single networks combining distance and phenotype predictions performed reasonably and were easier to use. In our testing, enlarging the output map of the FCNN from 1×1 to 3×3 pixels and adding ResNet-inspired skip connection appeared to be beneficial.

In summary, the final backbone of our system is a prediction of distance to cell center, which is designed for ease of annotation and post-processing (i.e. for a peak-finding algorithm), and reasonably approximates the shape of many lymphocytes, which are often round.

#### Network training, cell detection, and implementation

We used an Adam optimizer^52^ during training with a learning rate of 0.001, and mean squared error loss functions for both phenotype and distance pixel map predictions. Different weights were assigned to phenotype and distance map losses: 20 and 1 respectively. Phenotype loss functions could easily be replaced with categorical cross-entropy when binary predictions are desired, but this allowed us to both train on biologically more gradual markers like CD45RO, and include our Likert scale annotations in training. We normalize each channel per tile by using the default percentile-based normalization from the CSBdeep python library.^53^ During training we add Gaussian noise with a standard deviation of 0.1 to input and apply random data augmentations. Specifically, we perform horizontal and vertical flips and rotations, and change the input image intensities randomly in the same way as StarDist^14^ by first multiplying the input with a uniformly distributed scalar *s*1 *∼ U* (0.6, 2) and then adding another uniformly distributed scalar *s*2 *∼ U* (*−*0.2, 0.2). Convolutional layers perform batch normalization during training; fully connected layers perform dropout with a rate of 0.2.

We predict a circle with a radius of 5 pixels (2.5µm) around each annotated lymphocyte, with a value of 5 at the center, that drops to a value of 0 at the border of the circle. For pixels not part of annotated cells, we set the distance transform to -2. For each phenotype pseudomarker, the 5-point Likert scale is mapped to the values 0,0.25,0.5,0.75,1. To detect and phenotype cells based on the distance and pseudomarker predictions of the network, we first post-process the distance pixel map prediction with a Laplacian-of-Gaussian filter as implemented in scikit-image,^54^ using parameters min_sigma=3, max_sigma=5, and threshold=0.07. Then, for each detected cell location and each pseudomarker channel, we determine the mean expression value of that channel within a radius of 2 pixels around the center, and use this as the pseudomarker expression value of the cell.

The final network used in this paper was trained on 183,678 63*×*63*×*7 input environments taken from 27,888 annotations (each environment is centered near an annotated location, such that each annotation is covered by multiple environments). Training was run for 100 epochs, which took about 12 hours. Neural networks were constructed with TensorFlow,^55^ with important post-processing done in NumPy,^56^ SciPy^57^ and scikit-image.^54^

### Tumor and stroma tissue segmentation

Tumor and stroma tissue were identified using adaptive thresholding of different fluorescent stainings, as implemented by scikit-image.^54^ For tumor segmentation the tumor marker staining was thresholded, for stroma a sum of all stainings was thresholded to first get general tissue area, and identified tumor area was subtracted to get all non-tumor tissue. Tissue specific values were manually selected for tumor and stroma thresholds. Accurately segmenting tumor or stromal tissue is not trivial, but neural network methods based on U-net have shown success at tackling this task. While this simple segmentation algorithm likely does not achieve optimal performance, it is intended as *clear box* alternative to inForm software’s tissue segmentation. Replacing inForm with our own tissue segmentation in this manner allows us to present an tumor image analysis pipeline that is fully transparent and inspectable. Yet, our segmentation finds tumor and tissue in comparable regions to inForm software (**Supplementary Figure 3)**.

## Data Availability

A subset of the immunohistochemistry images used to train the model (AgarCyto data) and the corresponding annotations are publicly available for download at https://zenodo.org/record/5638697; this data can be used to test the provided model training and inference scripts (see below). The remaining training data and annotations are available upon request and will be made publicly available upon publication of this manuscript.

## Code Availability

The source code of the ImmuNet model, including scripts to train the model and to perform inference, are available at our GitHub repository at https://github.com/jtextor/immunet under the MIT license. The model is implemented in Python 3.6.9 and Tensorflow 1.14.0. The final trained model is available at https://zenodo.org/record/5638697.

## Acknowledgements

JT and SS were supported by the Dutch Cancer Society - Alpe d’HuZes foundation (grant 10620). JT and FB were supported by NWO grant VI.Vidi.192.084. MG, LvdW, KV, and CF were supported by Dutch Cancer Society grants 10673 and 2017-8244. CF was also supported by a grant from Oncode Institute and by ERC advanced grant ARTimmune 834618. EM was supported by a grant from the Hanarth Fonds (to JT).

We thank Dr. Willem Vreuls and Dr. Roland Pennings for providing tonsil tissues.

## Author contributions statement

SS and JT designed the machine learning framework. MG, LvdW, SvW, and KV provided critical feedback during development of the framework that led to substantial improvements. SS and JT and wrote software for data processing, quality control, network training, validation, visualization, and data export with important contributions from EM. FB contributed important empirical analyses during the development phase of the framework. SS, MG, EM, LvdW, FB, SvW, KV, and JT annotated cells to train the network and performed quality control. SS, JT and EM performed accuracy analyses. SS performed spatial statistics analyses and generated the CPM data.

## Competing interests statement

The authors declare no competing interests.

## Supplementary Tables

**Supplementary Figure 1:**
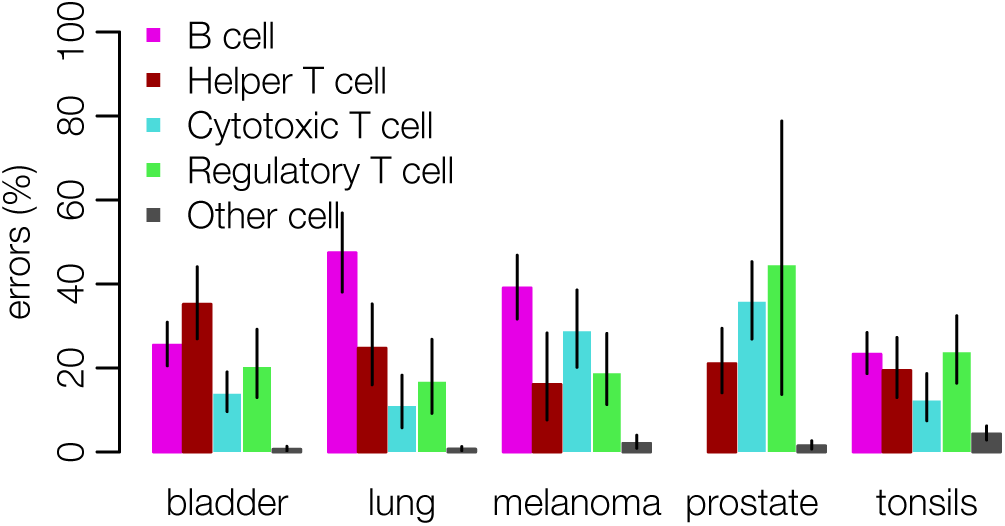
**Performance of inForm-based cell phenotyping** on the same data used in **Figure 3E**.

**Supplementary Figure 2:**
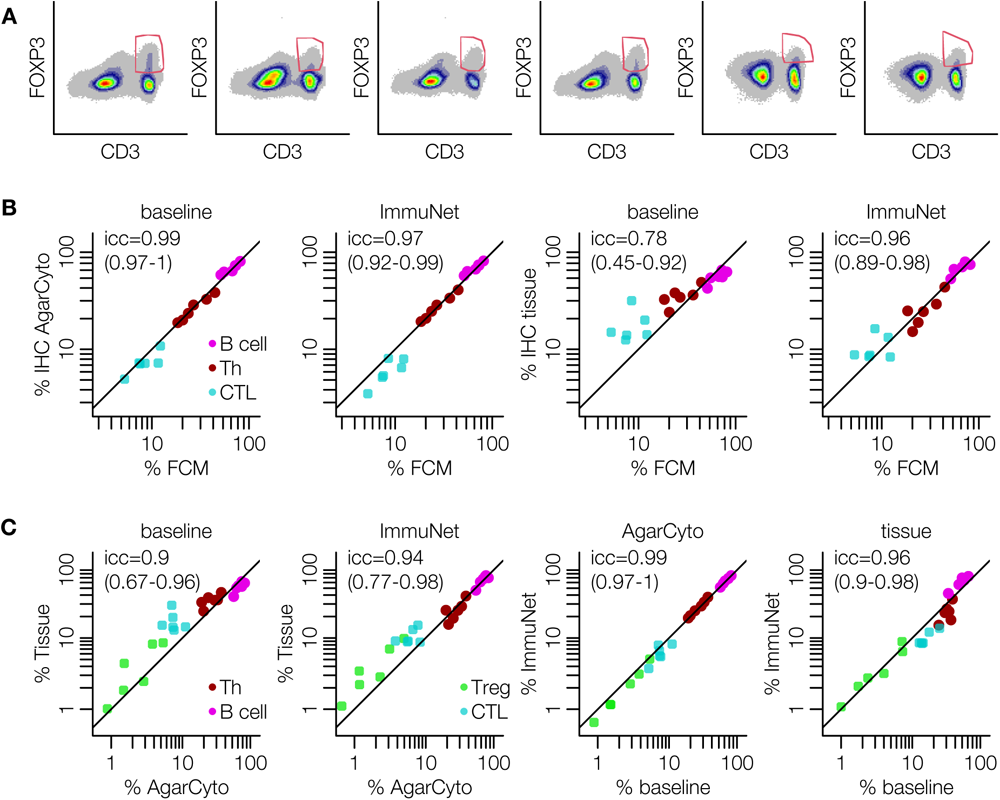
Influence of FOXP3 gating uncertainty on Treg quantification and comparison to mIHC data. **(A)** Gates used to identify FOXP3^+^ cells as a basis for Treg phenotyping. **(B)** Repetition of the analysis shown in **Figure 5** without Tregs. **(C)** Comparisons between mIHC measurements show higher agreement than to FACS, especially concerning the Treg population.

**Supplementary Figure 3:**
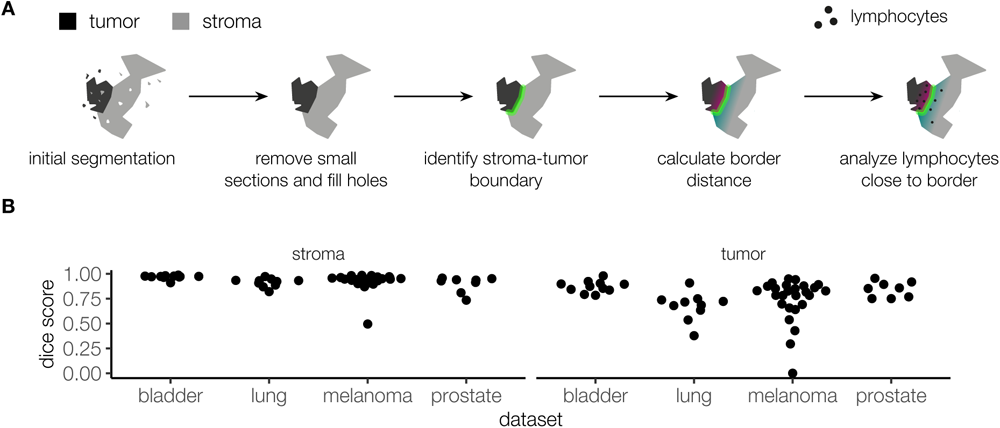
Stroma and tumor segmentation in whole-slide images. **(A)** To identify tumor and stroma tissue regions, we use adaptive thresholding with post-processing (see Methods). The border between tumor and non-tumor tissue is identified, and lymphocytes up to 100μm from border (infiltrated or outside) are selected for downstream analysis. **(B)** To compare our stroma and tumor tissue segmentation to that of inForm, we applied the fragment removal and hole filling steps of our algorithm to the output of inForm’s tissue segmentation. Subsequently, we compared the two segmentations using a pixel-wise Dice score. Boxplots show the distribution of Dice scores across all whole-slide images from each dataset.

**Supplementary Table 1:**
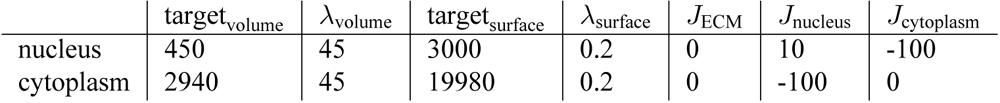
Cellular Potts model settings.

| | $\text{target}_{\text{volume}}$ | $\lambda_{\text{volume}}$ | $\text{target}_{\text{surface}}$ | $\lambda_{\text{surface}}$ | $J_{\text{ECM}}$ | $J_{\text{nucleus}}$ | $J_{\text{cytoplasm}}$ |
| --- | --- | --- | --- | --- | --- | --- | --- |
| nucleus | 450 | 45 | 3000 | 0.2 | 0 | 10 | -100 |
| cytoplasm | 2940 | 45 | 19980 | 0.2 | 0 | -100 | 0 |

**Supplementary Table 2:** The detailed ImmuNet architecture. In all convolutional and max pooling layers stride 1 is used.

| Layer | Filter dimension | Dilation rate | Output size | Connected to |
| --- | --- | --- | --- | --- |
| **Block 1** |  |  |  |  |
| **Main branch** |  |  |  |  |
| Convolution (conv1) | 64 x 7 x 4 x 4 | 1 | 60 x 60 x 64 | input |
| Batch normalization (bn1) |  |  |  | conv1 |
| ReLu activation (relu1) |  |  |  | bn1 |
| Convolution (conv2) | 64 x 64 x 3 x 3 | 1 | 58 x 58 x 64 | relu1 |
| Batch normalization (bn2) |  |  |  | conv2 |
| **Skip connection branch** |  |  |  |  |
| Convolution (skip_conv1) | 64 x 6 x 6 x 6 | 1 | 58 x 58 x 64 | input |
| Add (add1) |  |  |  | bn2, skip_conv1 |
| ReLu activation (relu2) |  |  |  | add1 |
| Max pooling (max1) | 2 x 2 | 1 | 57 x 57 x 64 | relu2 |
| **Block 2** |  |  |  |  |
| **Main branch** |  |  |  |  |
| Convolution (conv3) | 128 x 64 x 3 x 3 | 2 | 53 x 53 x 128 | max1 |
| Batch normalization (bn3) |  |  |  | conv3 |
| ReLu activation (relu3) |  |  |  | relu3 |
| Convolution (conv4) | 128 x 128 x 3 x 3 | 2 | 49 x 49 x 128 | relu3 |
| Batch normalization (bn4) |  |  |  | conv4 |
| **Skip connection branch** |  |  |  |  |
| Convolution (skip_conv2) | 128 x 64 x 5 x 5 | 2 | 49 x 49 x 128 | max1 |
| Add (add2) |  |  |  | bn4, skip_conv2 |
| ReLu activation (relu4) |  |  |  | add2 |
| Max pooling (max2) | 2 x 2 | 2 | 47 x 47 x 128 | relu4 |
| **Block 3** |  |  |  |  |
| **Main branch** |  |  |  |  |
| Convolution (conv5) | 256 x 128 x 3 x 3 | 4 | 39 x 39 x 256 | max2 |
| Batch normalization (bn5) |  |  |  | conv5 |
| ReLu activation (relu5) |  |  |  | bn5 |
| Convolution (conv6) | 256 x 256 x 3 x 3 | 4 | 31 x 31 x 256 | relu5 |
| Batch normalization (bn6) |  |  |  | conv6 |
| **Skip connection branch** |  |  |  |  |
| Convolution (skip_conv3) | 256 x 128 x 5 x 5 | 4 | 31 x 31 x 256 | max2 |
| Add (add3) |  |  |  | bn6, skip_conv3 |
| ReLu activation (relu6) |  |  |  | add3 |
| Max pooling (max3) | 2 x 2 | 4 | 27 x 27 x 256 | relu6 |
| Convolution (conv7) | 512 x 256 x 4 x 4 | 8 | 3 x 3 x 512 | max3 |
| Batch normalization (bn7) |  |  |  | conv7 |
| ReLu activation (relu7) |  |  |  | bn7 |
| Convolution (conv8) | 512 x 512 x 1 x 1 | 1 | 3 x 3 x 512 | relu7 |
| Dropout 0.2 (dp1) |  |  |  | conv8 |
| **Distance branch** |  |  |  |  |
| Convolution (conv9) | 512 x 512 x 1 x 1 | 1 | 3 x 3 x 512 | dp1 |
| ReLu activation (relu8) |  |  |  | conv9 |
| Dropout 0.2 (dp2) |  |  |  | relu8 |
| Convolution (conv10) | 1 x 512 x 1 x 1 | 1 | 3 x 3 x 1 | dp2 |
| Linear activation (linear1) |  |  |  | conv10 |
| **Phenotyping branch** |  |  |  |  |
| Convolution (conv11) | 512 x 512 x 1 x 1 | 1 | 3 x 3 x 512 | dp1 |
| ReLu activation (relu9) |  |  |  | conv11 |
| Dropout 0.2 (dp3) |  |  |  | relu9 |
| Convolution (conv12) | 5 x 512 x 1 x 1 | 1 | 3 x 3 x 5 | dp3 |
| Linear activation (linear2) |  |  |  | conv12 |

## Notes

https://github.com/jtextor/immunet

https://zenodo.org/record/5638697

### Competing Interest Statement

The authors have declared no competing interest.

### Summary of Updates

Edited text throughout for clarity and language; Generated more manually annotated data to evaluate the model; Improved the baseline model to ensure fairer comparison; Figures 2-5 revised; New Figure 6 added; New Supplementary Figures 1-2, 3a added (old Supplementary Figure 1 is now Supplementary Figure 3b); Author affiliations updated.

## References

1. van der Laak J, Litjens G, and Ciompi F. Deep learning in histopathology: the path to the clinic. Nature Medicine, 27(5):775–784, 2021. doi:10.1038/s41591-021-01343-4.

2 Dong C and Martinez GJ. T cells: the usual subsets. https://www.nature.com/documents/nri_posters_tcellsubsets.pdf. Accessed: 2021-10-18.

3 Gorris MAJ, Halilovic A, Rabold K, van Duffelen A, Wickramasinghe IN, Verweij D, Wortel IMN, Textor JC, de Vries IJM, and Figdor CG. Eight-color multiplex immunohistochemistry for simultaneous detection of multiple immune checkpoint molecules within the tumor microenvironment. The Journal of Immunology, 200(1):347–354, 2017. doi:10.4049/jimmunol.1701262.

4 Goltsev Y, Samusik N, Kennedy-Darling J, Bhate S, Hale M, Vazquez G, Black S, and Nolan GP. Deep profiling of mouse splenic architecture with CODEX multiplexed imaging. Cell, 174(4):968–981.e15, 2018. doi:10.1016/j.cell.2018.07.010.

5 Bjornson ZB, Nolan GP, and Fantl WJ. Single-cell mass cytometry for analysis of immune system functional states. Current Opinion in Immunology, 25(4):484–494, 2013. doi:10.1016/j.coi.2013.07.004.

6 Danaher P, Kim Y, Nelson B, Griswold M, Yang Z, Piazza E, and Beechem JM. Advances in mixed cell deconvolution enable quantification of cell types in spatially-resolved gene expression data. bioRxiv doi:10.1101/2020.08.04.235168, 2020.

7 Tan WCC, Nerurkar SN, Cai HY, Ng HHM, Wu D, Wee YTF, Lim JCT, Yeong J, and Lim TKH. Overview of multiplex immunohistochemistry/immunofluorescence techniques in the era of cancer immunotherapy. Cancer Communications, 40(4):135–153, 2020. doi:10.1002/cac2.12023.

8 Valen DAV, Kudo T, Lane KM, Macklin DN, Quach NT, DeFelice MM, Maayan I, Tanouchi Y, Ashley EA, and Covert MW. Deep learning automates the quantitative analysis of individual cells in live-cell imaging experiments. PLOS Computational Biology, 12(11):e1005177, 2016. doi:10.1371/journal.pcbi.1005177.

9 Berryman S, Matthews K, Lee JH, Duffy SP, and Ma H. Image-based phenotyping of disaggregated cells using deep learning. Communications Biology, 3(1), 2020. doi:10.1038/s42003-020-01399-x.

10. Ma J, Xie R, Gupta A, Almeida J, Bader GD, and Wang B, editors. Proceedings of The Cell Segmentation Challenge in Multi-modality High-Resolution Microscopy Images, volume 212 of Proceedings of Machine Learning Research. PMLR, 2023.

11. Park J, Choi W, Tiesmeyer S, Long B, Borm LE, Garren E, Nguyen TN, Tasic B, Codeluppi S, Graf T, Schlesner M, Stegle O, Eils R, and Ishaque N. Cell segmentation-free inference of cell types from in situ transcriptomics data. bioRxiv doi:10.1101/800748, 2020.

12. Meijering E. Cell segmentation: 50 years down the road [life sciences]. IEEE Signal Processing Magazine, 29(5):140–145, 2012. doi:10.1109/msp.2012.2204190.

13. Abdolhoseini M, Kluge MG, Walker FR, and Johnson SJ. Segmentation of heavily clustered nuclei from histopathological images. Scientific Reports, 9(1), 2019. doi:10.1038/s41598-019-38813-2.

14. Schmidt U, Weigert M, Broaddus C, and Myers G. Cell detection with star-convex polygons. In Medical Image Computing and Computer Assisted Intervention - MICCAI 2018 - 21st International Conference, Granada, Spain, September 16-20, 2018, Proceedings, Part II, pages 265–273, 2018. doi:10.1007/978-3-030-00934-2_30.

15. Weigert M, Schmidt U, Haase R, Sugawara K, and Myers G. Star-convex polyhedra for 3d object detection and segmentation in microscopy. In The IEEE Winter Conference on Applications of Computer Vision (WACV), 2020. doi:10.1109/WACV45572.2020.9093435.

16. Graner F and Glazier JA. Simulation of biological cell sorting using a two-dimensional extended potts model. Physical Review Letters, 69(13):2013–2016, 1992. doi:10.1103/physrevlett.69.2013.

17. Savill NJ and Hogeweg P. Modelling morphogenesis: From single cells to crawling slugs. Journal of Theoretical Biology, 184(3):229–235, 1997. doi:10.1006/jtbi.1996.0237.

18. Wortel IMN and Textor J. Artistoo, a library to build, share, and explore simulations of cells and tissues in the web browser. eLife, 10, 2021. doi:10.7554/elife.61288.

19. Stack EC, Wang C, Roman KA, and Hoyt CC. Multiplexed immunohistochemistry, imaging, and quantitation: A review, with an assessment of tyramide signal amplification, multispectral imaging and multiplex analysis. Methods, 70(1):46–58, 2014. doi:10.1016/j.ymeth.2014.08.016.

20. Vasaturo A, Halilovic A, Bol KF, Verweij DI, Blokx WAM, Punt CJA, Groenen PJTA, van Krieken JHJM, Textor J, de Vries IJM, and Figdor CG. T-cell landscape in a primary melanoma predicts the survival of patients with metastatic disease after their treatment with dendritic cell vaccines. Cancer Research, 76(12):3496–3506, 2016. doi:10.1158/0008-5472.can-15-3211.

21. Roelofsen T, Wefers C, Gorris MAJ, Textor JC, Massuger LFAG, de Vries IJM, and van Altena AM. Spontaneous regression of ovarian carcinoma after septic peritonitis; a unique case report. Frontiers in Oncology, 8, 2018. doi:10.3389/fonc.2018.00562.

22. van Beek JJ, Flórez-Grau G, Gorris MA, Mathan TS, Schreibelt G, Bol KF, Textor J, and de Vries IJM. Human pDCs are superior to cDC2s in attracting cytolytic lymphocytes in melanoma patients receiving DC vaccination. Cell Reports, 30(4):1027–1038.e4, 2020. doi:10.1016/j.celrep.2019.12.096.

23. Boudewijns S, Bloemendal M, de Haas N, Westdorp H, Bol KF, Schreibelt G, Aarntzen EHJG, Lesterhuis WJ, Gorris MAJ, Croockewit A, van der Woude LL, van Rossum MM, Welzen M, de Goede A, Hato SV, van der Graaf WTA, Punt CJA, Koornstra RHT, Gerritsen WR, Figdor CG, and de Vries IJM. Autologous monocyte-derived DC vaccination combined with cisplatin in stage III and IV melanoma patients: a prospective, randomized phase 2 trial. *Cancer Immunology*, Immunotherapy, 69(3):477–488, 2020. doi:10.1007/s00262-019-02466-x.

24. Hoeijmakers YM, Gorris MA, Sweep FC, Bulten J, Eysbouts YK, Massuger LF, Ottevanger PB, and de Vries JI. Immune cell composition in the endometrium of patients with a complete molar pregnancy: Effects on outcome. Gynecologic Oncology, 160(2):450–456, 2021. doi:10.1016/j.ygyno.2020.11.005.

25. Rodriguez-Rosales YA, Langereis JD, Gorris MA, van den Reek JM, Fasse E, Netea MG, de Vries IJM, Gomez-Muñoz L, van Cranenbroek B, Körber A, Sondermann W, Joosten I, de Jong EM, and Koenen HJ. Immunomodulatory aged neutrophils are augmented in blood and skin of psoriasis patients. Journal of Allergy and Clinical Immunology, 148(4):1030–1040, 2021. doi:10.1016/j.jaci.2021.02.041.

26. PerkinElmer Inc. inForm user manual. https://www.perkinelmer.com/Content/LST_Software_Downloads/in-FormUserManual_2_3_0_rev1.pdf.

27. Wang W, Taft DA, Chen YJ, Zhang J, Wallace CT, Xu M, Watkins SC, and Xing J. Learn to segment single cells with deep distance estimator and deep cell detector. Computers in Biology and Medicine, 108:133–141, 2019. doi:10.1016/j.compbiomed.2019.04.006.

28. Hopcroft JE and Karp RM. An n^5/2 algorithm for maximum matchings in bipartite graphs. SIAM Journal on computing, 2(4):225–231, 1973.

29. Kerstens HM, Robben JC, Poddighe PJ, Melchers WJ, Boonstra H, de Wilde PC, Macville MV, and Hanselaar AG. AgarCyto: A novel cell-processing method for multiple molecular diagnostic analyses of the uterine cervix. Journal of Histochemistry & Cytochemistry, 48(5):709–718, 2000. doi:10.1177/002215540004800515.

30. Oble DA, Loewe R, Yu P, and Mihm J Martin C. Focus on tils: prognostic significance of tumor infiltrating lymphocytes in human melanoma. Cancer immunity, 9:3–3, 2009.

31. Tumeh PC, Harview CL, Yearley JH, Shintaku IP, Taylor EJM, Robert L, Chmielowski B, Spasic M, Henry G, Ciobanu V, West AN, Carmona M, Kivork C, Seja E, Cherry G, Gutierrez AJ, Grogan TR, Mateus C, Tomasic G, Glaspy JA, Emerson RO, Robins H, Pierce RH, Elashoff DA, Robert C, and Ribas A. PD-1 blockade induces responses by inhibiting adaptive immune resistance. Nature, 515(7528):568–571, 2014. doi:10.1038/nature13954.

32. Liu X, Wu S, Yang Y, Zhao M, Zhu G, and Hou Z. The prognostic landscape of tumor-infiltrating immune cell and immunomodulators in lung cancer. Biomedicine & Pharmacotherapy, 95:55–61, 2017. doi:10.1016/j.biopha.2017.08.003.

33. Huh JW, Lee JH, and Kim HR. Prognostic Significance of Tumor-Infiltrating Lymphocytes for Patients With Colorectal Cancer. Archives of Surgery, 147(4):366–372, 2012. doi:10.1001/archsurg.2012.35.

34. van der Woude LL, Gorris MA, Halilovic A, Figdor CG, and de Vries IJM. Migrating into the tumor: a roadmap for t cells. Trends in Cancer, 3(11):797–808, 2017. doi:10.1016/j.trecan.2017.09.006.

35. Balkenhol MC, Ciompi F, Świderska-Chadaj Ż, van de Loo R, Intezar M, Otte-Höller I, Geijs D, Lotz J, Weiss N, de Bel T, Litjens G, Bult P, and van der Laak JA. Optimized tumour infiltrating lymphocyte assessment for triple negative breast cancer prognostics. The Breast, 56:78–87, 2021. doi:10.1016/j.breast.2021.02.007.

36. Schapiro D, Jackson HW, Raghuraman S, Fischer JR, Zanotelli VRT, Schulz D, Giesen C, Catena R, Varga Z, and Bodenmiller B. histocat: analysis of cell phenotypes and interactions in multiplex image cytometry data. Nature Methods, 14(9):873–876, 2017. doi:10.1038/nmeth.4391.

37. McQuin C, Goodman A, Chernyshev V, Kamentsky L, Cimini BA, Karhohs KW, Doan M, Ding L, Rafelski SM, Thirstrup D, Wiegraebe W, Singh S, Becker T, Caicedo JC, and Carpenter AE. CellProfiler 3.0: Next-generation image processing for biology. PLOS Biology, 16(7):e2005970, 2018. doi:10.1371/journal.pbio.2005970.

38. Berg S, Kutra D, Kroeger T, Straehle CN, Kausler BX, Haubold C, Schiegg M, Ales J, Beier T, Rudy M, Eren K, Cervantes JI, Xu B, Beuttenmueller F, Wolny A, Zhang C, Koethe U, Hamprecht FA, and Kreshuk A. ilastik: interactive machine learning for (bio)image analysis. Nature Methods, 2019. doi:10.1038/s41592-019-0582-9.

39. Kensert A, Harrison PJ, and Spjuth O. Transfer learning with deep convolutional neural networks for classifying cellular morphological changes. SLAS DISCOVERY: Advancing the Science of Drug Discovery, 24(4):466–475, 2019. doi:10.1177/2472555218818756.

40. Kim S, Baek J, Park J, Kim G, and Kim S. Instaformer: Instance-aware image-to-image translation with transformer. In Proceedings of the IEEE/CVF Conference on Computer Vision and Pattern Recognition (CVPR), pages 18321–18331, 2022.

41. Gorris MAJ, van der Woude LL, Kroeze LI, Bol K, Verrijp K, Amir AL, Meek J, Textor J, Figdor CG, and de Vries IJM. Paired primary and metastatic lesions of patients with ipilimumab-treated melanoma: high variation in lymphocyte infiltration and hla-abc expression whereas tumor mutational load is similar and correlates with clinical outcome. Journal for ImmunoTherapy of Cancer, 10(5):e004329, 2022. doi:10.1136/jitc-2021-004329.

42. van der Woude LL, Gorris MAJ, Wortel IMN, Creemers JHA, Verrijp K, Monkhorst K, Grünberg K, van den Heuvel MM, Textor J, Figdor CG, Piet B, Theelen WSME, and de Vries IJM. Tumor microenvironment shows an immunological abscopal effect in patients with nsclc treated with pembrolizumab-radiotherapy combination. Journal for ImmunoTherapy of Cancer, 10(10):e005248, 2022. doi:10.1136/jitc-2022-005248.

43. van Wilpe S, Simnica D, Slootbeek P, van Ee T, Pamidimarri Naga S, Gorris MAJ, van der Woude LL, Sultan S, Koornstra RHT, van Oort IM, Gerritsen WR, Kroeze LI, Simons M, van Leenders GJLH, Binder M, de Vries IJM, and Mehra N. Homologous recombination repair deficient prostate cancer represents an immunologically distinct subtype. OncoImmunology, 11(1), 2022. doi:10.1080/2162402x.2022.2094133.

44. van Wilpe S, Gorris MAJ, van der Woude LL, Sultan S, Koornstra RHT, van der Heijden AG, Gerritsen WR, Simons M, de Vries IJM, and Mehra N. Spatial and temporal heterogeneity of tumor-infiltrating lymphocytes in advanced urothelial cancer. Frontiers in Immunology, 12, 2022. doi:10.3389/fimmu.2021.802877.

45. Graham Martínez C, Barella Y, Kus Öztürk S, Ansems M, Gorris MA, van Vliet S, Marijnen CA, and Nagtegaal ID. The immune microenvironment landscape shows treatment-specific differences in rectal cancer patients. Frontiers in Immunology, 13, 2022. doi:10.3389/fimmu.2022.1011498.

46. Theelen WSME, Peulen HMU, Lalezari F, van der Noort V, de Vries JF, Aerts JGJV, Dumoulin DW, Bahce I, Niemeijer ALN, de Langen AJ, Monkhorst K, and Baas P. Effect of pembrolizumab after stereotactic body radiotherapy vs pembrolizumab alone on tumor response in patients with advanced non–small cell lung cancer. JAMA Oncology, 5(9):1276, 2019. doi:10.1001/jamaoncol.2019.1478.

47. Moen E, Bannon D, Kudo T, Graf W, Covert M, and Van Valen D. Deep learning for cellular image analysis. Nature Methods, 16(12):1233–1246, 2019. doi:10.1038/s41592-019-0403-1.

48. Shelhamer E, Long J, and Darrell T. Fully convolutional networks for semantic segmentation. arxiv:1605.06211, https://arxiv.org/abs/1605.06211, 2016.

49. Ronneberger O, Fischer P, and Brox T. U-net: Convolutional networks for biomedical image segmentation. arxiv:1505.04597 https://arxiv.org/abs/1505.04597, 2015.

50. Swiderska-Chadaj Z, Pinckaers H, van Rijthoven M, Balkenhol M, Melnikova M, Geessink O, Manson Q, Sherman M, Polonia A, Parry J, Abubakar M, Litjens G, van der Laak J, and Ciompi F. Learning to detect lymphocytes in immunohistochemistry with deep learning. Medical Image Analysis, 58:101547, 2019. doi:https://doi.org/10.1016/j.media.2019.101547.

51. Hermsen M, Volk V, Bräsen JH, Geijs DJ, Gwinner W, Kers J, Linmans J, Schaadt NS, Schmitz J, Steenbergen EJ, Swiderska-Chadaj Z, Smeets B, Hilbrands LB, Feuerhake F, and van der Laak JAWM. Quantitative assessment of inflammatory infiltrates in kidney transplant biopsies using multiplex tyramide signal amplification and deep learning. Laboratory Investigation, 2021. doi:10.1038/s41374-021-00601-w.

52. Kingma DP and Ba J. Adam: A method for stochastic optimization. arxiv: 1412.6980, https://arxiv.org/abs/1412.6980, 2017.

53. Weigert M, Schmidt U, Boothe T, Müller A, Dibrov A, Jain A, Wilhelm B, Schmidt D, Broaddus C, Culley S, Rocha-Martins M, Segovia-Miranda F, Norden C, Henriques R, Zerial M, Solimena M, Rink J, Tomancak P, Royer L, Jug F, and Myers EW. Content-aware image restoration: pushing the limits of fluorescence microscopy. Nature Methods, 15(12):1090–1097, 2018. doi:10.1038/s41592-018-0216-7.

54. van der Walt S, Schönberger JL, Nunez-Iglesias J, Boulogne F, Warner JD, Yager N, Gouillart E, Yu T, and the scikit-image contributors. scikit-image: image processing in Python. PeerJ, 2:e453, 2014. doi:10.7717/peerj.453.

55. Abadi M, Agarwal A, Barham P, Brevdo E, Chen Z, Citro C, Corrado GS, Davis A, Dean J, Devin M, Ghemawat S, Goodfellow I, Harp A, Irving G, Isard M, Jia Y, Jozefowicz R, Kaiser L, Kudlur M, Levenberg J, Mané D, Monga R, Moore S, Murray D, Olah C, Schuster M, Shlens J, Steiner B, Sutskever I, Talwar K, Tucker P, Vanhoucke V, Vasudevan V, Viégas F, Vinyals O, Warden P, Wattenberg M, Wicke M, Yu Y, and Zheng X. TensorFlow: Large-scale machine learning on heterogeneous systems, 2015. Software available from tensorflow.org.

56. Harris CR, Millman KJ, van der Walt SJ, Gommers R, Virtanen P, Cournapeau D, Wieser E, Taylor J, Berg S, Smith NJ, Kern R, Picus M, Hoyer S, van Kerkwijk MH, Brett M, Haldane A, del Río JF, Wiebe M, Peterson P, Gérard-Marchant P, Sheppard K, Reddy T, Weckesser W, Abbasi H, Gohlke C, and Oliphant TE. Array programming with NumPy. Nature, 585(7825):357–362, 2020. doi:10.1038/s41586-020-2649-2.

57. Virtanen P, Gommers R, Oliphant TE, Haberland M, Reddy T, Cournapeau D, Burovski E, Peterson P, Weckesser W, Bright J, van der Walt SJ, Brett M, Wilson J, Millman KJ, Mayorov N, Nelson ARJ, Jones E, Kern R, Larson E, Carey CJ, Polat İ, Feng Y, Moore EW, VanderPlas J, Laxalde D, Perktold J, Cimrman R, Henriksen I, Quintero EA, Harris CR, Archibald AM, Ribeiro AH, Pedregosa F, van Mulbregt P, and SciPy 1.0 Contributors. SciPy 1.0: Fundamental Algorithms for Scientific Computing in Python. Nature Methods, 17:261–272, 2020. doi:10.1038/s41592-019-0686-2.

